# Aclarubicin stimulates RNA polymerase II elongation at closely spaced divergent promoters

**DOI:** 10.1101/2023.01.09.523323

**Authors:** Matthew Wooten, Brittany Takushi, Kami Ahmad, Steven Henikoff

## Abstract

Anthracyclines are a class of widely prescribed anti-cancer drugs that disrupt chromatin by intercalating into DNA and enhancing nucleosome turnover. To understand the molecular consequences of anthracycline-mediated chromatin disruption, we utilized CUT&Tag to profile RNA polymerase II during anthracycline treatment in *Drosophila* cells. We observed that treatment with the anthracycline aclarubicin leads to elevated levels of elongating RNA polymerase II and changes in chromatin accessibility. We found that promoter proximity and orientation impacts chromatin changes during aclarubicin treatment, as closely spaced divergent promoter pairs show greater chromatin changes when compared to codirectionally-oriented tandem promoters. We also found that aclarubicin treatment changes the distribution of non-canonical DNA G-quadruplex structures both at promoters and at G-rich pericentromeric repeats. Our work suggests that the anti-cancer activity of aclarubicin is driven by the effects of nucleosome disruption on RNA polymerase II, chromatin accessibility and DNA structures.

## Introduction

Chromatin proteins shape the physical structure of DNA by regulating accessibility, torsion, and 3D association (1–3). Nucleosomes, the fundamental unit of eukaryotic chromatin structure, are comprised of two copies each of histones H3, H4, H2B and H2A and wrap ~1.7 turns, or 146 base pairs of DNA (4). Nucleosomes present a formidable barrier to transcription initiation and elongation, and numerous auxiliary factors are required to remove or remodel nucleosomes to allow for efficient transcriptional initiation and progression. RNA polymerase II (RNA Pol II) dynamics associated with active transcription can also substantially impact chromatin structure. Transcription can lead to dynamic changes including histone turnover, addition of post-transcriptional modification, and changes in histone variant composition (5). Transcription can also induce changes in DNA structure: unwound, or negatively supercoiled DNA generated in the wake of the advancing polymerase can promote the formation of non-canonical DNA structures, such as G-quadruplexes (2, 6–8), which are comprised of G-rich DNA held together by Hoogsteen base-pairing arrangements (9).

Changes in chromatin structure can also be induced by small molecule drugs that intercalate between DNA bases, such as anthracyclines, a class of anticancer drugs given to over 1 million patients per year (10). Most anthracyclines used in the clinic also poison topoisomerases (11), which are enzymes responsible for relieving torsional strain on DNA (12, 13). A popular model for the anti-cancer effect of anthracyclines is that by poisoning topoisomerase, anthracyclines generate DNA damage, leading to programmed cell death (11). However, this model has been challenged by studies demonstrating that anthracycline intercalation promotes nucleosome turnover and causes chromatin disruption (10, 14–19). One particular anthracycline – aclarubicin – generates high levels of histone eviction without causing DNA damage, while the anthracycline doxorubicin causes only moderate levels of histone eviction but high levels of DNA damage (16). These differences have potential clinical implications, as chromatin disruption is better correlated with cancer cell death than DNA damage (20). However, it remains unclear how extensive anthracycline-induced chromatin disruption is, how this may differ between anthracyclines, and what the downstream impacts of chromatin disruption are on cellular functions such as transcription.

In this study, we investigated the impacts of the anthracycline-mediated chromatin disruption on RNA Pol II in *Drosophila melanogaster* cells, a system where transcriptional regulation has been extensively characterized (21–25). We find that aclarubicin, but not doxorubicin, induces gains in both elongating RNA Pol II and chromatin accessibility. We also find that distinct genomic regions show differential responses to aclarubicin treatment: closely spaced divergent promoter elements show greater increases in RNA Pol II and chromatin accessibility when compared to more distantly spaced promoter elements. We also show that aclarubicin treatment induces chromatin disruption of the pericentromeric Dodeca-satellite repeat blocks, which display gains in RNA Pol II, chromatin accessibility and G-quadruplex formation following aclarubicin treatment. The distinct impacts that doxorubicin and aclarubicin have on chromatin suggest that the anti-cancer effects of these two anthracyclines are caused by distinct mechanisms.

## Results

### Aclarubicin treatment inhibits proliferation and increases elongating RNA Pol II in genes

To assess the impacts of drug treatment on cell morphology and viability, *Drosophila* Kc167 cells were resuspended in media containing 1μM doxorubicin, 1μM aclarubicin or 5 μM actinomycin D, an intercalating drug known to arrest RNA pol II elongation in gene bodies (Fig. 1A). After 24 hours of drug treatment, all three drugs similarly inhibited cell proliferation compared to mock treated (DMSO) controls (Fig. 1B). Previous studies have shown that anthracyclines can also impact nucleolar structure (26). Using antibodies to fibrillarin as a nucleolar marker, we observed nucleolar signal in 97% of doxorubicin-treated cells after 24 hours treatment, demonstrating that at this dose the nucleolus is not disrupted. In contrast, only 15% of aclarubicin-treated cells had nucleolar fibrillarin signal, indicating disruption of rDNA transcription in most cells (fig. S1A, D). Thus, aclarubicin is a more effective disruptor of nucleolar structure than doxorubicin, despite these drugs’ structural similarities. Similarly, only 11% of actinomycin D-treated cells show nucleoli, indicating potent disruption of rDNA transcription. Using antibodies to cleaved caspase as a marker of apoptosis, we observed that after 24 hours treatment, both aclarubicin- and actinomycin D-treated cells showed high levels of cell death compared to controls, whereas doxorubicin-treated cells showed little increase (fig. S1B, E). Finally, we assessed cell size after drug treatment. Cells treated with doxorubicin were slightly larger than control cells, whereas both aclarubicin- and actinomycin D-treated cells were smaller than control cells (fig. S1C). Measurements of nucleolar integrity, size, and death after 1 hour of drug treatment showed no significant changes for any of three drugs (fig. S1A-C). Therefore, we performed all subsequent experiments after 30 minutes of drug exposure, prior to observable signs of toxicity, to assess immediate effects of drug treatment on chromatin.

**Fig. 1:**
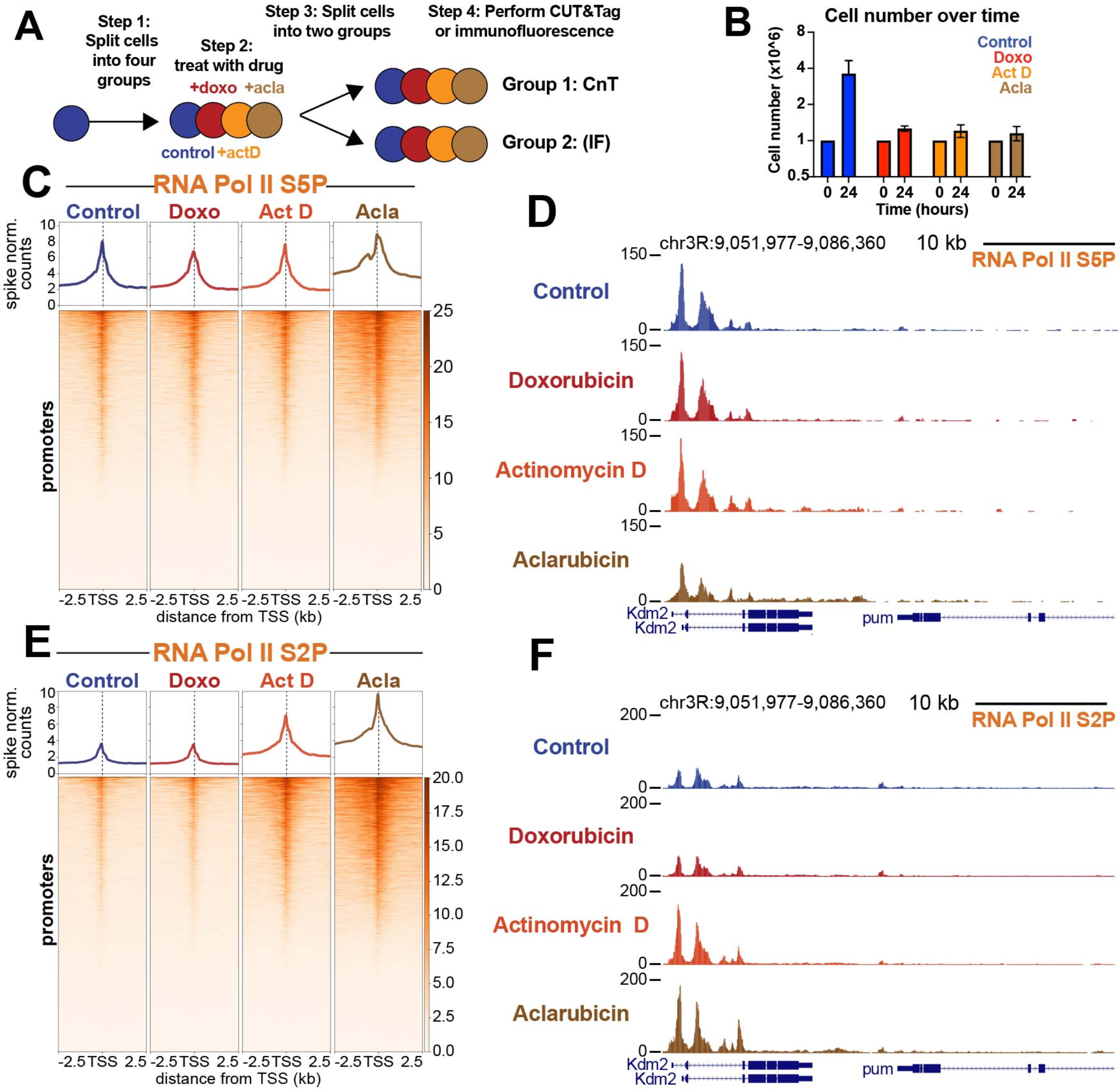
Aclarubicin blocks growth and causes gains in RNA pol II ser2P. (**A**) Experimental design. The four treatments were 1μM Doxorubicin (doxo), 5μM actinomycin D (Act D)1μM aclarubicin (acla) and mock-treated (DMSO) control (**B**) Cell numbers at 0 hour and at 24 hours. (**C**) Heatmap aligned to TSS of all promoters showing paused polymerase spike-normalized signal under each treatment condition. (**D**) Representative UCSC browser track snapshot of paused RNA Pol II distribution. (**E**) Heatmap aligned to the transcriptional start site (TSS) of all promoters showing spike-normalized elongating polymerase signal under each treatment condition. (**F**) Representative UCSC browser track snapshot of elongating polymerase distribution.

Nucleosomes can block RNA Pol II access to DNA at promoters (27–29). Therefore, active promoters are remodeled to remove nucleosomes and facilitate assembly of the transcriptional pre-initiation complex (PIC) (30–32). As previous work has shown that anthracyclines stimulate nucleosome turnover at gene promoters, we sought to understand whether anthracycline treatment affects RNA Pol II dynamics. To assess changes in RNA Pol II, we treated cells with drugs for 30 minutes and then performed <u>Cl</u>eavage <u>U</u>nder <u>T</u>argets and <u>Tag</u>mentation (CUT&Tag) (33, 34) targeting both paused RNA Pol II (marked by Serine-5 phosphorylation (Ser5P)) and elongating RNA Pol II (marked by Serine-2 phosphorylation (Ser2P)) (35). Importantly, all CUT&Tag experiments profiling RNA Pol II were quantified using a human cell post-treatment spike-in that enables direct comparisons of chromatin factor levels between different samples treated across a single experiment (34, 36). To visualize the amounts and distribution of RNA Pol II, we generated spike-normalized coverage heatmaps at all 16,972 promoters in the *Drosophila melanogaster* genome centered on their transcription start site (TSS). While paused RNA Pol II-Ser5P showed minimal changes across all three drugs (Fig. 1C, D), aclarubicin-treated samples gained large amounts of elongating RNA Pol II-Ser2P at and around active promoters (Fig. 1E, F). Samples treated with actinomycin D also gained a modest amount of elongating RNA Pol II signal, similar to previous observations (37, 38). In contrast, elongating RNA Pol II-Ser2P signal in doxorubicin-treated samples was unaffected (Fig. 2E, F). The observation that aclarubicin but not doxorubicin increases RNA Pol II elongation reveals a fundamental difference in the mechanisms underlying chromatin disruption in these distinct anthracyclines.

**Fig. 2:**
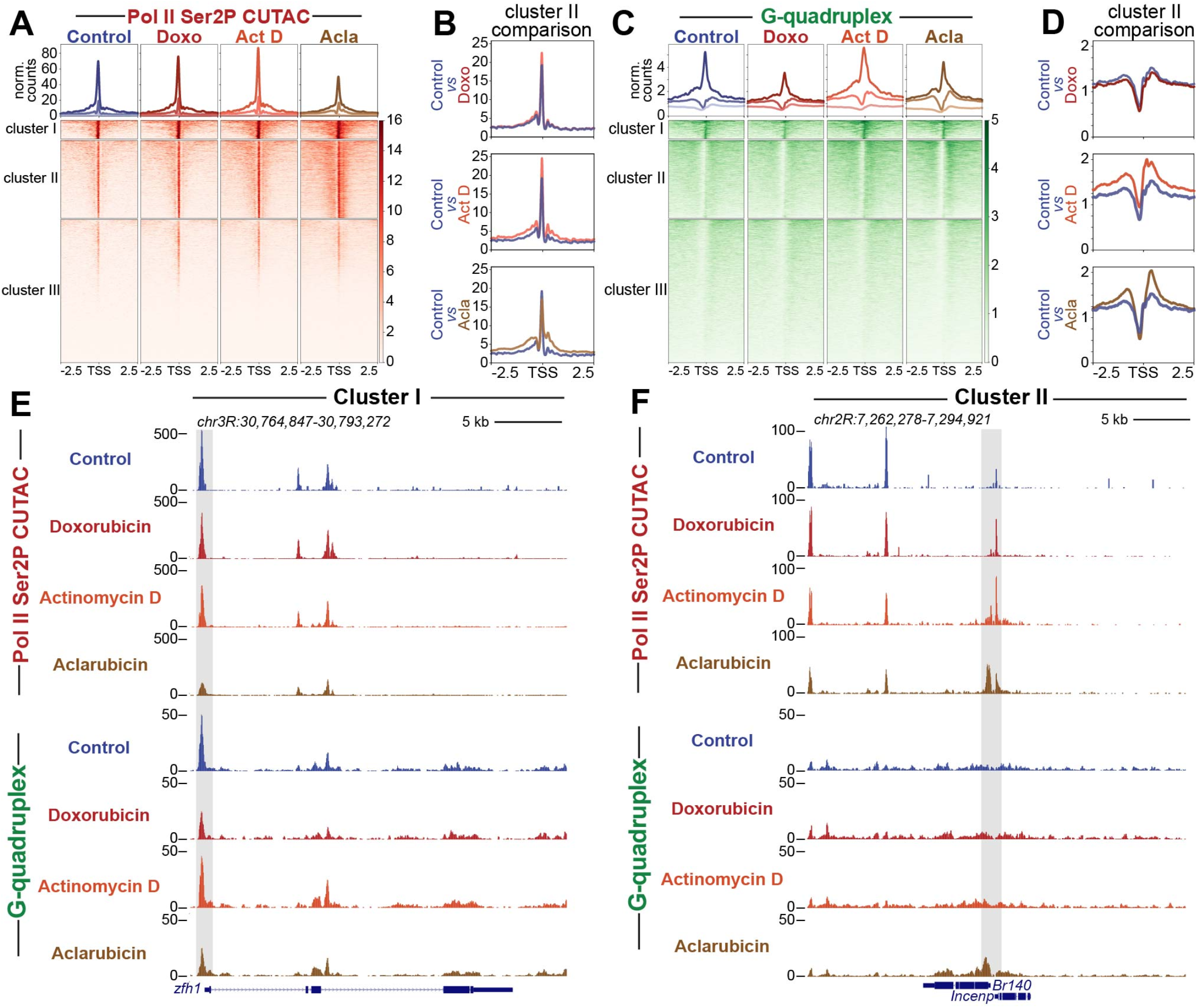
Drug treatment impacts chromatin accessibility and G-quadruplex formation. (**A**) Heatmap aligned to TSS of all promoters showing normalized counts of CUTAC chromatin accessibility targeting RNA Pol II ser2P clustered via k-means clustering (k=3). (**B**) Enlarged comparison of accessibility differences between different drug groups and controls. (**C**) Heatmap aligned to the transcriptional start site (TSS) of all promoters showing normalized counts of G-quadruplex CUT&Tag signal clustered via k-means clustering (k=3). (**D**) Enlarged comparison of G-quadruplex differences between different drug groups and controls. (**E**) Representative UCSC browser track snapshot of CUTAC and G-quadruplex distribution at cluster I gene. Gray box indicates promoter region. (**F**) Representative UCSC browser track snapshot of G-quadruplex distribution at Cluster II gene. Gray box indicates promoter region.

### Aclarubicin promotes chromatin accessibility and G-quadruplex formation

We next wondered whether the increase in RNA Pol II caused by aclarubicin might affect chromatin accessibility and G-quadruplex formation, both of which have been associated with gene activation (39, 40). To assay chromatin accessibility changes during drug treatment, we treated cells with each drug for thirty minutes and then performed <u>C</u>leavage <u>U</u>nder <u>T</u>argeted <u>A</u>ccessible <u>C</u>hromatin (CUTAC) (33). CUTAC uses low-salt tagmentation conditions while tethering pA-Tn5 to RNA Pol II to produce chromatin accessibility maps with low background. To assess changes in G-quadruplexes during drug treatment, we performed CUT&Tag using the BG4 antibody, which binds specifically to G-quadruplex structures (41). Importantly, CUT&Tag does not require fixation or sonication, making it ideal for profiling G-quadruplexes which may be disrupted by such harsh treatments. Previous studies have demonstrated that CUT&Tag using the BG4 antibody is effective in profiling G-quadruplex structures (42, 43).

To visualize changes in chromatin accessibility during drug treatment, we performed k-means clustering of *Drosophila* promoters, and generated heatmaps to compare accessibility across different groups (Fig. 2A). In control conditions, Clusters I, II and III are distinguished by levels of chromatin accessibility: Cluster I (n = 1306) promoters are highly accessible, Cluster II (n = 5517) promoters are moderately accessible, and Cluster III (n = 10149) promoters display background levels of accessibility. Doxorubicin-treated samples showed minimal changes across all clusters, while actinomycin D-treated samples showed subtle gains in accessibility at regions proximal to the TSS (Fig 2A, B, E, F). Aclarubicin-treated samples, by contrast, showed reduced accessibility at Cluster I promoters, and increased accessibility upstream and downstream of the TSS at Cluster II promoters (Fig. 2A, B, E, F).

We then used the same clusters generated by CUTAC data to assess changes in G-quadruplex formation between drug treatments. Doxorubicin-treated samples showed a loss of G-quadruplex signal at Cluster I promoters, and no change in G-quadruplex signal at Cluster II. Aclarubicin-treated samples showed a loss of G-quadruplex signal at Cluster I, but gains in G-quadruplex signal at Cluster II promoters. Actinomycin D showed subtle gains in G-quadruplex signal across all promoter clusters, similar to previous reports demonstrating that actinomycin D can increase G-quadruplex formation at promoters (43) (Fig. 2C, D, E, F). Interestingly, regions showing the greatest gain in G-quadruplex signal during aclarubicin treatment were downstream of the TSS, distinct from the G-quadruplexes in Cluster I that are located upstream of the TSS. The distinct spatial distribution of G-quadruplexes in Cluster I and Cluster II, and their contrasting responses to aclarubicin treatment suggest that these G-quadruplexes may be formed and/or regulated in distinct ways.

Lastly, to assess changes in RNA Pol II at distinct gene clusters, we sorted RNA Pol II-Ser5P and RNA Pol II-Ser2P datasets using k-means clusters generated by CUTAC data. Similar to trends observed in CUTAC and G-quadruplex datasets, we observed that aclarubicin-treated samples showed gains in RNA Pol II-Ser2P and RNA Pol II-Ser5P at Cluster II promoters (fig. S2A-D, 2F). These gains in RNA Pol II signal were observable upstream and downstream of the promoter region (fig. S2B and 2D). Actinomycin D-treated samples showed subtle gains in Cluster I and Cluster II, whereas doxorubicin-treated samples showed minimal changes across all clusters (fig. S2A-F). Together, these data demonstrate that drug treatment differentially impacts the distribution of RNA Pol II on distinct groups of genes.

### Closely spaced divergent promoters show greater chromatin disruption than distant promoters

We sought to understand what features differentiate genes in Cluster I from genes in Cluster II. While examining gene promoters in each cluster, we noticed that Cluster I promoters are frequently isolated in the genome, whereas Cluster II promoters are frequently found close to gene neighbors. To assess whether proximity to neighboring genes correlates with differential responsiveness to anthracyclines, we sorted promoters by their distance to the nearest upstream promoter and generated heatmaps of chromatin features. Indeed, closely spaced promoters in drug-treated samples appear to show greater gains in chromatin accessibility, G-quadruplex formation, and RNA Pol II-Ser2P/5P than more distant promoters (Fig. 3A, fig. S3A-C).

**Fig. 3:**
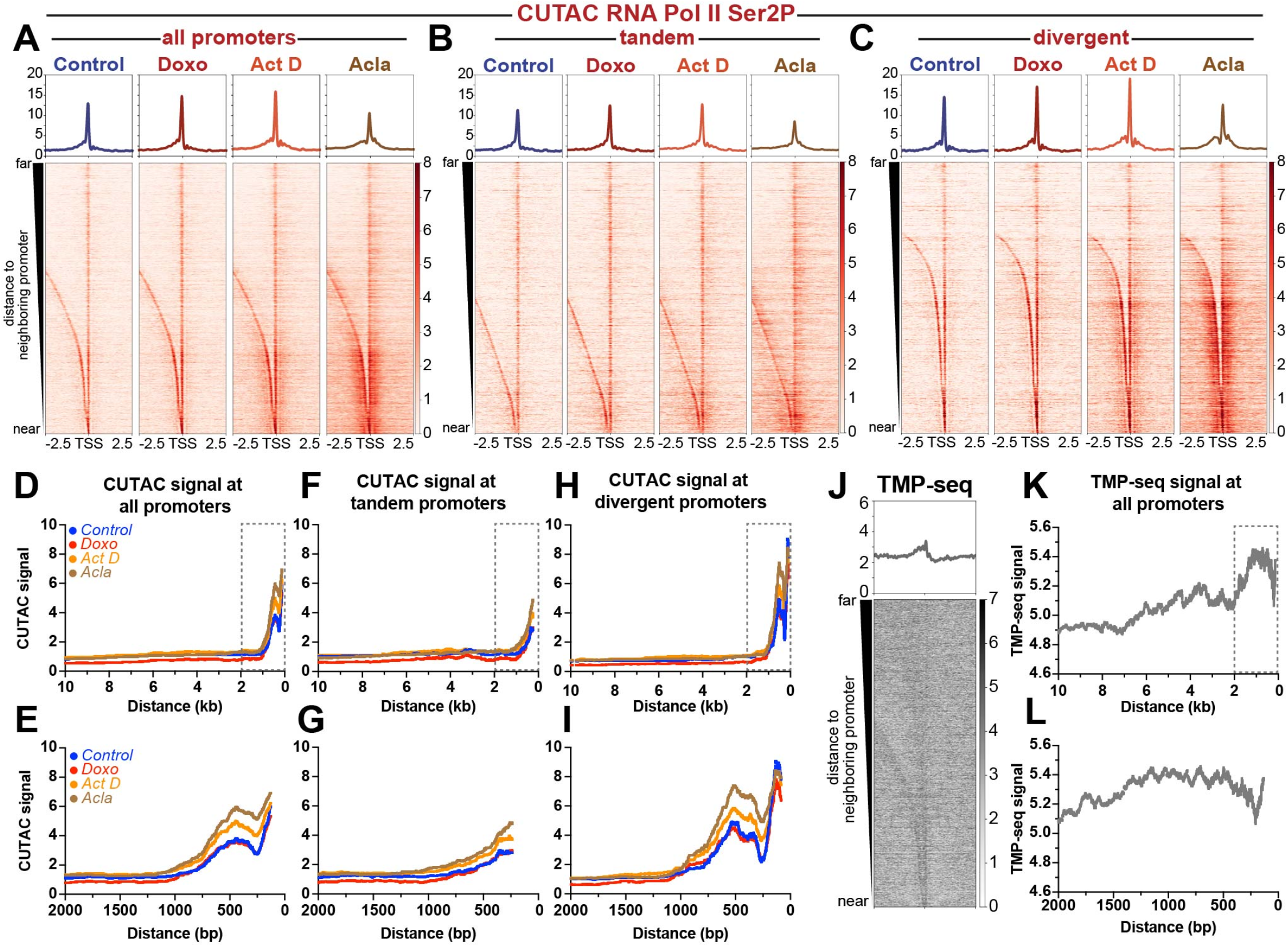
Promoter distance and orientation effects chromatin accessibility and superhelical torsion. Heatmaps of normalized counts showing CUTAC chromatin accessibility signal targeting RNA Pol II ser2P aligned to the transcriptional start site (TSS) of all promoters (**A**), tandem promoters (**B**) and divergent promoters (**C**) sorted by distance to nearest upstream promoter element and plotted in descending order. (**D**) Plot showing moving median of CUTAC chromatin accessibility as a function of distance between neighboring promoter elements for all promoters. (**E**) Zoom in of boxed region in panel D. (**F**) Plot showing moving median of CUTAC chromatin accessibility as a function of distance between neighboring promoter elements for tandem promoters. (**G**) Zoom in of boxed region in panel F. (H) Plot showing moving median of CUTAC chromatin accessibility as a function of distance between neighboring promoter elements for all divergent promoters. (**I**) Zoom in of boxed region in panel H. (**J**) Heatmap of tri-methyl psoralen-seq (TMP-seq) data aligned to the transcriptional start site (TSS) of all promoters, sorted by distance to nearest upstream promoter element and plotted in descending order. (**K**) Plot showing moving average of TMP-seq data as a function of distance between neighboring promoter elements for all promoters. (**L**) Zoom in of boxed region in panel K.

To better assess the quantitative differences in accessibility between closely spaced *versus* distant promoters, we plotted CUTAC signal for each treatment as a function of distance between promoters. All treatment groups showed increased accessibility at closely spaced promoter elements when compared to distantly spaced promoter elements (Fig. 3D, E). Doxorubicin-treated samples showed minimal overall differences in accessibility when compared to control samples (Fig. 3D, E). Quantification of accessibility for promoters with an inter-promoter distance less than 2kb revealed a slight, but statistically significant difference in accessibility for doxorubicin-treated samples (median = 2.8) when compared to controls (median = 2.6; fig. S3D). For promoters spaced greater than 2kb apart, doxorubicin-treated samples showed lower accessibility values (median = 0.6) when compared to controls (median = 1.0; fig. S3D). Actinomycin D-treated samples showed no statistically-significant difference for promoters greater than 2kb apart (median = 1.1 act D *versus* median = 1.0 control), and greater accessibility for promoters less than 2kb apart (median = 3.6 act D *versus* median = 2.6 control; Fig. 3D, E, Fig. S3D). Aclarubicin-treated samples showed the most striking differences in accessibility as a function of distance. While distantly spaced promoters showed no difference in accessibility when compared to control samples, closely spaced promoters showed significantly greater median accessibility relative to controls. This switch in accessibility appears to occur at ~1kb (Fig. 3D, E), where aclarubicin-treated samples first begin to show consistently higher accessibility. Quantification of median accessibility for closely spaced promoters (>2kb) confirmed that aclarubicin-treated samples show significantly higher accessibility (median = 4.3 acla *versus* median = 2.6 control), while distantly spaced promoters show no significant difference (median = 1.0 acla *versus* median = 1.0 control; fig. S3D). These results demonstrate that aclarubicin-treatment disrupts chromatin structure most effectively at closely spaced promoters.

As closely spaced promoters show the greatest changes in chromatin structure during drug treatment, we wondered whether these promoters are transcribed in the same (tandem) or divergent directions. We found that 58% (5372 out of 9288) of the promoters less than 2kb apart are divergently oriented, with that number growing to 68% (4604 out of 6788) for promoters positioned closer than 1kb. To determine whether accessibility differed in closely spaced divergent promoters *versus* closely spaced tandem promoters, we separated these two groups of promoters and plotted the accessibility changes during drug treatment (Fig. 3B, C, 3F-I.) We observed that divergent promoters showed greater levels of accessibility overall and greater increases in accessibility following drug treatment when compared to tandem (co-directionally oriented) promoters (Fig. 3B, C. 3F-I). To further assess differences between divergent and tandem promoters across drug treatments, we plotted fold change in accessibility surrounding the TSS (fig. S3E-G) for each drug treatment. While doxorubicin-treated samples showed minimal differences in accessibility when compared to controls (fig. S3E), actinomycin D-treated samples showed modest increases in accessibility, with divergent promoters showing greater gains in accessibility compared to tandem promoters (fig. S3F). Aclarubicin-treated samples showed greatest increases in accessibility relative to controls, with divergent promoters once again showing greatest fold change in accessibility (fig. S3G). Based on these observations, we conclude that closely spaced, divergent genes are more sensitive than tandem genes to changes in chromatin structure during drug treatment.

Previous work from our lab proposed that negative supercoiling (underwound DNA) generated in the wake of transcribing RNA Pol II facilitates intercalation of anthracyclines into the genome (10), leading to chromatin disruption (44, 45). As transcriptionally active promoters can propagate negative superhelical torsion over 1kb upstream of the TSS (28, 46, 47), promoters in close proximity may be affected by supercoiling generated at nearby promoters. To determine whether differences in negative supercoiling distinguish closely spaced promoters from isolated promoters, we re-analyzed trimethyl-psoralen sequencing data (TMP-seq) from Teves et al. (28), which profiled the incidence of negative supercoiling in the *Drosophila melanogaster* genome. We observed that closely spaced promoters showed higher levels of negative supercoiling when compared to distantly spaced promoters (Fig. 3J-L). As negative supercoiling should facilitate intercalation of anthracyclines (48, 49), concentrated negative supercoiling upstream of closely spaced divergent promoters may facilitate drug binding, resulting in chromatin disruption. As chromatin structure around promoters is critical for regulating RNA Pol II dynamics, it is likely that intercalation at these sites would be more impactful in driving changes in RNA Pol II elongation as well as gains in promoter proximal chromatin accessibility.

### Histone genes and pericentromeric Dodeca-satellite repeats show distinct responses to drug treatment

The histone locus cluster is a region of the genome with an especially high incidence of closely spaced divergent promoters (Fig. 4A). Given the elevated levels of chromatin disruption observed at closely spaced divergent promoter elements genome-wide, we wondered whether the histone locus would also show changes in chromatin structure during anthracycline treatment. We found that, similar to divergent promoters genomewide, the histone locus showed large gains in both accessibility (Fig. 4B) and elongating RNA Pol II (Ser2P) (Fig. 4C) following treatment with aclarubicin or actinomycin D. Interestingly, the histone locus did not show similar gains in initiating RNA Pol II-Ser5P or G-quadruplex formation (fig. S4A-B), further suggesting that the strongest immediate impacts of aclarubicin-mediated chromatin disruption are on elongating RNA Pol II. Since high levels of free histone proteins can induce cell death (50–52), this may contribute to the cytotoxicity of aclarubicin.

**Fig. 4:**
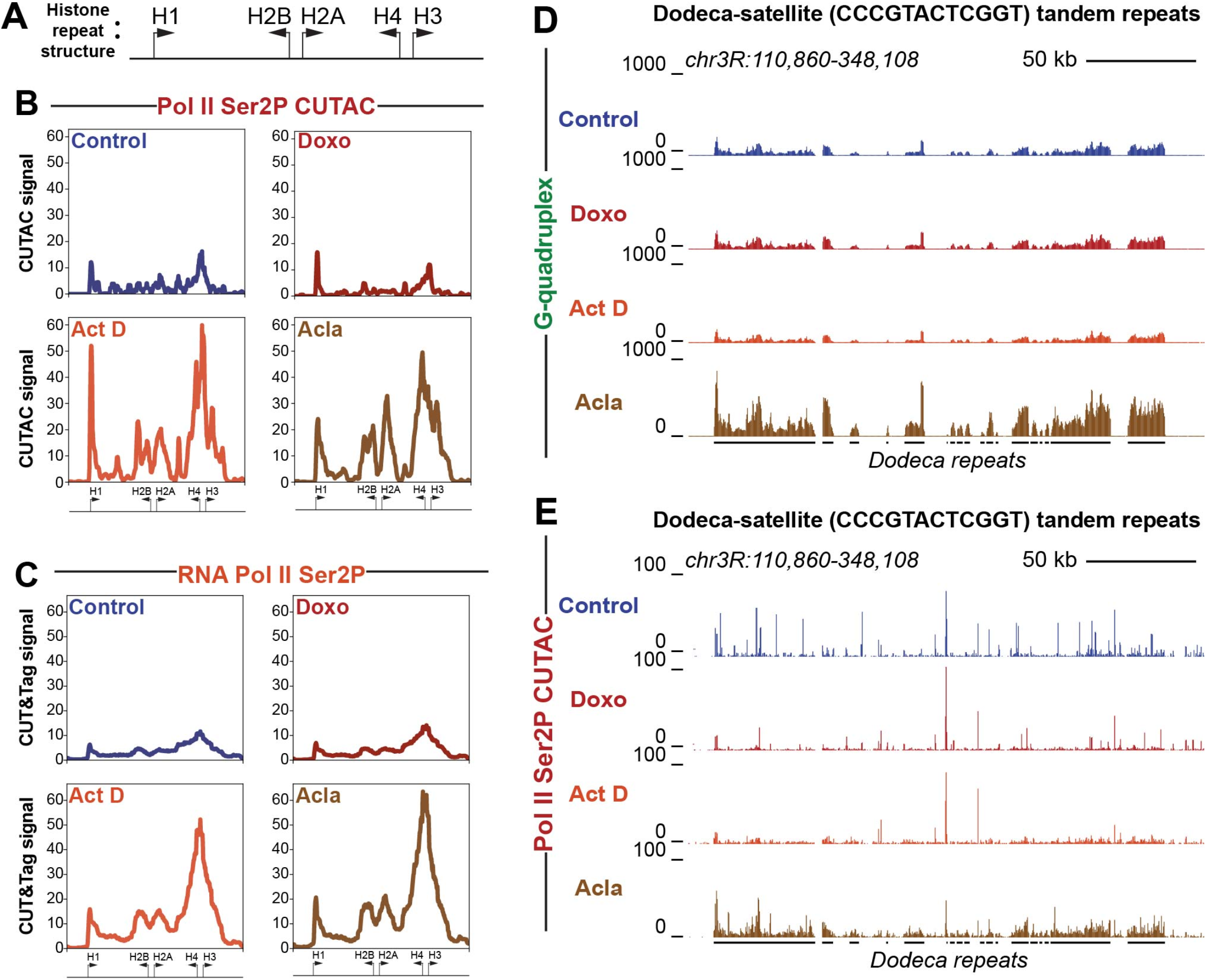
Histone cluster and Dodeca-satellite repeats show distinct responses to drug treatment. (**A**) Cartoon showing divergent orientation of histone cluster genes. Arrows indicate direction of transcription with the base of the arrow representing approximate promoter position. (**B**) Average coverage plot of histone cluster showing normalized counts of CUTAC signal targeting RNA Pol II ser2P. (**C**) Average coverage plot of histone clusters showing normalized counts of CUT&Tag data targeting RNA Pol II ser2P. Arrows at the bottom of average plots indicate approximate positions of histone genes cartooned in panel A. (**D**) UCSC browser track snapshot of G-quadruplex CUT&Tag data at Dodeca-satellite repeats. (**E**) UCSC browser track snapshot of RNA Pol II Ser2P CUTAC data at Dodeca-satellite repeats. Black lines below the browser tracks indicate location of Dodeca-satellite repeats.

Previous studies have shown that the Dodeca-satellite (CCCGTACTCGGT) tandem repeats, found exclusively in a pericentromeric region *of Drosophila melanogaster* chromosome 3, can form non-canonical structures known as i-motifs from the repetitive runs of cytosines found within the repeat (53). We wondered if G-quadruplexes form on the opposite strand from i-motif structures at Dodeca-satellite and if they are affected by drug treatment. Indeed, we found that G-quadruplex signal is detectable at Dodeca-satellite repeats in control datasets (Fig. 4D). Strikingly, G-quadruplex formation increased in cells treated with aclarubicin. This effect did not occur in cells treated with doxorubicin or with actinomycin D. Aclarubicin treatment also induced chromatin accessibility at the Dodeca-satellite repeats (Fig. 4E), as well as gains in RNA Pol II-Ser2P and RNA Pol II-Ser5P (fig. S4C, D), while treatment with doxorubicin and actinomycin D did not. These data demonstrate that aclarubicin, but not doxorubicin or actinomycin D is able to disrupt chromatin structure at this tandemly repetitive region of the genome.

## Discussion

We have shown here that treatment of *Drosophila* cells with the anthracycline aclarubicin results in gains in RNA Pol II-Ser2P and changes in chromatin accessibility and G-quadruplex formation at active gene promoters. Closely spaced divergent promoter pairs are particularly sensitive to drug treatment, likely due to high levels of negative supercoiling enriched at these promoter elements. Chromatin effects are not limited to active gene promoters and can also be found at certain repetitive sequences prone to forming non-canonical DNA structures. Our observations reveal molecular features underlying anthracycline-mediated chromatin disruption and demonstrate clear differences between doxorubicin and aclarubicin treatment, suggesting that these drugs may exert anti-cancer effects through different mechanisms.

The predominant model for anthracycline cytotoxicity is that these drugs induce DNA damage by preventing topoisomerase II from ligating together the double-strand breaks made during topoisomerase-mediated DNA cleavage (54). However, anthracyclines also disrupt chromatin structure by evicting histones (16) and increasing nucleosome turnover around promoters (17). Chromatin disruption is better correlated to cell death than DNA damage, suggesting that chromatin disruption is the principle anti-cancer activity of anthracyclines (16, 20, 45). Our results here demonstrate that aclarubicin, which is permissive for topoisomerase II ligation of double-strand breaks, promotes RNA Pol II elongation and chromatin accessibility at active promoters. Aclarubicin treatment does not affect the frequency of RNA Pol II binding, implying that aclarubicin targets the conversion of initiating RNA pol II into the elongating form. This effect may occur through aclarubicin-mediated disruption of the first nucleosome downstream of an active promoter. The first (+1) nucleosome is critical in regulating RNA Pol II elongation rates (21, 29), and disruption of the +1 nucleosome would remove a barrier to polymerase elongation. Thus, chromatin disruption and nucleosome turnover induced by aclarubicin has widespread effects on the transcriptional status of cells.

Previous studies have shown that aclarubicin promotes higher levels histone turnover than doxorubicin (16). These observations have prompted investigations into what molecular features of anthracyclines are responsible for driving histone disruption. Anthracyclines are tetracyclic molecules with an anthraquinone backbone connected by a glycosidic linkage to a sugar moiety (20). Recent studies have found that N,N-dimethylation of the carbohydrate appended to the anthraquinone aglycon is critical for both chromatin disruption and cytotoxicity, and the authors of this study speculated that this modification alters the interaction dynamics of anthracyclines with DNA (14). As previous studies have shown that doxorubicin-DNA aminal adducts can form between the 3’-NH2 of the doxorubicin sugar, the N2 of the guanine base, and formaldehyde (55, 56), the addition of two methyl groups (as in the case of aclarubicin) to the critical amino sugar might convert these drugs from a covalent DNA intercalator into a reversible DNA intercalator, affecting the dynamics by which these drugs perturb the contacts between DNA and the surface of the nucleosome. While it remains unclear what specific molecular features are responsible for aclarubicin’s distinct effect on chromatin structure, these studies emphasized that chemical modification that increase chromatin damage are highly coordinated with cancer cell death whereas chemical modifications associated with increased DNA damage are not.

It is striking that the chromatin effects of aclarubicin treatment are particularly severe at closely spaced divergent promoters. We show that these promoters are enriched for negative supercoiling, likely due transcriptional activity of promoter pairs (28, 46, 47). As negative supercoiling facilitates intercalation of small molecules such as anthracyclines into DNA (48, 49), we speculate that increased negative supercoiling between closely spaced divergent promoters could drive greater drug binding, leading to greater chromatin damage. Interestingly, a recent study in patient breast cancer samples and cell lines revealed that elevated expression of factors that promote chromatin accessibility and gene expression, such as COMPASS, BAF, KDM4B and KAT6B, lead to increased sensitivity to anthracycline treatment, whereas factors that promote chromatin compaction, such as PRC2, lead to anthracycline resistance (57). These studies suggest that the chromatin environment could be playing a critical role in rendering cancer cells more susceptible to the influence of anthracyclines, possibly by increasing the accessibility of anthracyclines to DNA. These observations are in agreement with previous studies (58), which have proposed that instability in cancer cell chromatin (58–60) could be a factor in rendering cancer cells more susceptible to the impacts of intercalating drugs when compared to healthy cells (58).

The effects of aclarubicin are not limited to active gene promoters, as aclarubicin treatment is able to generate elevated levels of both RNA Pol II-Ser2P and RNA Pol II-Ser5P at Dodeca-satellite repeats. These sequences are not normally transcribed in somatic cells but can form non-canonical DNA structures (53). Drug intercalation may trigger misfolding of these DNAs, stimulating their transcription (61). Our results imply that aclarubicin broadly alters chromosome structure, which may contribute to its cytotoxic properties.

Interestingly, other chemotherapeutic agents such as curaxins have also been shown to intercalate into DNA, drive nucleosome turnover, and induce aberrant DNA structures (58, 62). Like anthracyclines, the amount of chromatin disruption caused by curaxins is correlated with cytotoxicity, whereas DNA damage is not (62). Thus, chromatin disruption may be a major mechanism of action for distinct chemotherapies. Such a model is encouraging, as DNA damage contributes to many undesirable off-target effects of anthracyclines, including cardiotoxicity and therapy-related tumors (16, 63).

## Methods

### Cell culture and drug treatment

*Drosophila* Kc167 cells grown to log phase in HYQ-SFX Insect medium (Invitrogen) supplemented with 18 mM L-Glutamine and harvested as previously described (28). All cell counts and measures of cell size were measured using the Vi-CELL XR Cell Viability Analyzer (https://www.beckman.com). Doxorubicin (Sigma-Aldrich; D1515-10MG), aclarubicin (Cayman Chemical Company; 15993), and actinomycin D (Sigma-Aldrich; A9415-2MG) were resuspended to 10 mM in DMSO and frozen in aliquots. For cell treatments, compounds were added to cell medium containing 1.5 x 10^6^ cells per mL to a final concentration of 1 μM (aclarubicin, doxorubicin) or 5 μM (actinomycin D) and incubated at room temperature (RT). Cells were harvested after 30 minutes for imaging and CUT&Tag profiling, or for 1 hour or 24 hours to assess growth rates and cell size, cell death and nucleolus structure.

### Immunofluorescent labelling

Samples of 500,000 Kc167 cells were harvested, spun down and resuspended in ice cold PBS with 0.1% Triton and 0.1% formaldehyde and incubated for 10 minutes on ice. Cells were spun down and resuspended in ice cold PBS with 0.1% Triton-X100. Cells were then spun down onto a clean glass slide (Fisherbrand Superfrost Plus Microscope Slides; ThermoFisher Scientific, cat. no. 12-550-15) using a ThermoScientific Shandon Cytospin 4. Slides were washed twice with 50 mL 1× PBS each time and placed in a humid chamber with 1 mL of blocking solution (2.5% BSA in 1× PBST) for 30 minutes of pre-blocking. Blocking buffer was then drained, and primary antibodies were added for incubation overnight at 4 °C. Slides were then washed twice with 50 mL 1× PBS and incubated with secondary antibodies for 2 hours at RT. Slides were then washed twice with 50 mL 1× PBS and mounted with ProLong Diamond Mounting Media (Thermo Fisher Scientific: P36965). Cells were imaged on an EVOS auto 2.0 microscope (ThermoFisher).

### Whole-Cell CUT&Tag for chromatin profiling

Cleavage Under Targets and Tagmentation (CUT&Tag) was performed as described (64, 65) with some modifications. Briefly, 2 million *Drosophila* Kc167 cells were spun down and resuspended in 1 mL ice cold PBS containing 12,000 human K562 cells for spike-in normalization. Cells were spun down and resuspended in 1.5 mL wash buffer (20 mM HEPES at pH 7.5, 150 mM KCl, 0.5 mM spermidine and 1× protease inhibitors) by gentle pipetting. 5 μL of concanavalin A-coated magnetic beads (Smart-Lifesciences) were activated and added to cells and incubated for 10 minutes on ice. The supernatant was then removed, and bead-bound cells were resuspended in 100 μL dig-wash buffer (20 mM HEPES at pH 7.5, 150 mM KCl, 0.5 mM spermidine, 1× protease inhibitors, 0.05% digitonin) containing 2 mM EDTA and a 1:25 dilution of the 0.1 mg/mL BG4 primary antibody (Sigma-Aldrich: MABE917). Primary antibody incubation was performed overnight at 4°C and then the liquid was removed. Cells were washed 3x in 200 μL dig-wash buffer. Wash buffer was then drained, and cells were resuspended in 100 μL dig-wash buffer containing 2 mM EDTA and a 1:50 dilution of mouse anti-FLAG antibody (Sigma-Aldrich; F1804-1MG) and incubated at RT for 1 hour. Cells were washed 3x in 200 μL dig-wash buffer. Wash buffer was then drained, and cells were resuspended in 100 μL dig-wash buffer containing 2 mM EDTA and a 1:50 dilution of rabbit anti-mouse antibody (Sigma, M7023) and incubated at RT for 1 hour with slow rotation. A 1:50 dilution of pA-Tn5 adapter complex was prepared in dig-300 buffer (0.05% digitonin, 20 mM HEPES at pH 7.5, 300 mM KCl, 0.5 mM spermidine, 1× protease inhibitors). Fifty μL of pA-Tn5 adapter complex was then added to the cells with gentle vortexing, followed by incubation for 1 hour at RT. Cells were washed three times in 200 μL dig-300 buffer to remove unbound pA-Tn5 proteins. Cells were then immersed in 100 μL of tagmentation buffer (dig-300 buffer with 10 mM MgCl2) and incubated at 37°C for 1 hour. Cells were placed on a magnet and supernatant removed. Cells were washed with 50 μL 10 mM TAPS with 16.5 mM EDTA and resuspended in 100 μLs buffer containing 10 mM TAPS, 16.6mM EDTA, 0.1% SDS and 0.2 mg/mL proteinase K and incubated at 56°C for 1 hour. 200 μLs 300 wash buffer (20 mM HEPES at pH 7.5, 300 mM NaCl, 0.5 mM spermidine, 1× protease inhibitors) was then added to the tube, after which, the DNA was extracted via PCI for library preparation. Twenty-one μL DNA was mixed with a universal i5 and a uniquely barcoded i7 primer and amplified with NEB Q5 high-fidelity 2× master mix (cat# M0492S). The libraries were purified with 1.1 × volume of Sera-Mag carboxylate-modified magnetic beads and subjected to LabChip DNA analysis and Illumina sequencing. A similar methodology was used for profiling elongating RNA polymerase II (antiRNAPII-Ser2-phosphorylation antibody Cell Signaling Technology cat# 13499S) and initiating RNA polymerase II (anti-RNAPII-Ser5-phosphorylation antibody Cell Signaling Technology cat#13523s) using 80,000 *Drosophila melanogaster* whole cells. In some experiments, nuclei were extracted and profiled according the CUT&Tag-direct protocol (33). CUTAC was performed as described using an antibody to RNA polymerase II-Ser2-phosphorylation (66).

### CUT&Tag data processing and analysis

Libraries were sequenced on an Illumina HiSeq instrument with paired end 50×50 reads. Adapters were clipped by cutadapt (http://dx.doi.org/10.14806/ej.17.1.200) version 2.9 with parameters: -j 8 --nextseq-trim 20 -m 20 -a AGATCGGAAGAGCACACGTCTGAACTCCAGTCA -A AGATCGGAAGAGCGTCGTGTAGGGAAAGAGTGT -Z. Clipped reads were aligned by Bowtie2 (67) to the UCSC *Drosophila melanogaster* Dm6 reference sequence (68) with parameters: --very-sensitive-local --soft-clipped-unmapped-tlen --dovetail --no-mixed --no-discordant -q --phred33 -I 10 -X 1000. Clipped reads were also aligned by Bowtie2 (67) to the UCSC Homo sapiens HG19 reference sequence (68) with parameters: --end-to-end --very-sensitive --no-overlap --no-dovetail --no-mixed --no-discordant -q -- phred33 -I 10 -X 1000. Properly paired reads were extracted from the alignments by samtools (Version 1.9) (69). Spike-in calibrated *D. melanogaster* tracks in bigwig format were made by bedtools (70) 2.30.0 genomecov with a scaling factor of (10,000/number of fragments mapped to Homo sapiens). Normalized count tracks in bigwig format were also made by bedtools (70) 2.30.0 genomecov which are the fraction of counts at each base pair scaled by the size of the reference sequence (137,567,484) so that if the scaled counts were uniformly distributed there would be 1 at each position. Spike-in calibrated bigwig files were then uploaded to Galaxy (71) and heatmaps were generated using the computematrix function in deeptools (Version 3.5.1) (72). A bedfile containing a list of all *D. melanogaster* promoters was used by computematrix function to compare spike-normalized reads from bedgraph files aligned at all *D. melanogaster* promoters. The output of the computematrix function was visualized using the plotHeatmap function in Galaxy.

For k-means clustering analysis, we used the plotHeatmap function in Galaxy with k=3 on RNA Pol II-Ser2P CUTAC data. We then used this sorting for other datasets (G-quadruplex, RNA Pol II-Ser2P CUT&Tag and RNA Pol II-Ser5P CUT&Tag) by using the bedfile output of the plotHeapmap function as the “regions to plot” file for subsequent computeMatrix operations.

For analysis of nearby promoters, inter-promoter distance was calculated using promoter coordinates from the eukaryotic promoter database (https://epd.epfl.ch//index.php) (73). For each promoter in the database, the distance to the nearest promoter upstream was used to assign a distance value (in base pairs) to each promoter in the database. The strand information for each promoter was used to classify the nearest upstream promoter as tandem (same strand) or divergent (opposite strand). Promoters with the nearest promoter on the opposite strand were classified as divergent, whereas promoters with the nearest promoter on the same strand were classified as tandem.

To generate plots showing changes in CUTAC signal as a function of distance between promoters (Fig. 3D-I), total CUTAC signal at each promoter from 100 base pairs upstream to 500 base pairs downstream were summed using the multiBigwigSummary function from Galaxy deeptools. These values were calculated for all promoters across control, doxorubicin-treated, actinomycin D-treated, and aclarubicin-treated conditions. Promoters were then sorted by inter-promoter distance from greatest to least. A moving median value was then calculated to assess changes in accessibility with decreasing inter-promoter distances. We chose to calculate the median accessibility for each decile, or tenth of promoters with closest inter-promoter distances. As there are 16,972 promoters in the epd promoter database, 1697 represents 1/10^th^, or a decile of all promoters. Therefore, for each promoter we calculated the median CUTAC values for 1697 promoters with closest inter-promoter distances (848 promoters with greater inter-promoter distance, 848 promoters with lesser inter-promoter distance). We performed a similar calculation on the distance between promoters to give a moving median value of inter-promoter distance for each data point plotted on the graph. We then plotted the median CUTAC values as a function of the median distance. To generate plots showing TMP-seq as a function of distance, we calculated total TMP-seq signal 200 basepairs upstream of the TSS. This region was chosen, as this specifically looks at negative supercoiling upstream of the TSS at promoter regions. We then performed a similar moving average using deciles as described above. Two values were removed from decile analysis as they were extreme outliers: tx_1 and MOEH_1 had TMP-seq values greater than 64 standard deviations above the mean (mean = 5.21, stdev = 5.25 tx_1 = 846.79; MOEH_1 = 344.57).

To generate plots shown in fig. S3E-G comparing divergent and tandem promoters in control samples to divergent and tandem promoters in drug treated samples, the matrix values underlying the heatmaps in Fig. 3A and 3B were exported from galaxy using the computeMatrix function. These matrices contained normalized counts for each promoter broken up into 10 base pair (bp) bins. We then generated average normalized counts values for each drug treatment group and for each promoter orientation (divergent *versus* tandem) for every 10bp bind up to 2500bp surrounding the TSS. Average values at each 10bp bin were then divided (drug value/control value) to generate a fold-difference ratio plot surrounding the TSS.

## Acknowledgments

We thank our Fred Hutchinson Cancer Center colleagues, T. Llagas and D. Xu for help with cell culture, C. Codomo and T. Bryson for sequencing library pooling, and J. Henikoff for preparing the sequencing data for analysis.

## Funding

Howard Hughes Medical Institute (SH), Washington University Genome Training Grant T32HG000035 (MW)

## Author contributions

Conceptualization: MW, KA, SH

Methodology: MW, KA, SH

Investigation: MW, BT

Visualization: MW

Supervision: KA, SH

Writing—original draft: MW

Writing—review & editing: MW, KA, SH

## Competing interests

The authors declare that they have no competing interests.

## Data and materials availability

All data needed to evaluate the conclusions in the paper are present in the paper and/or the Supplementary Materials.

---

**Fig. S1:**
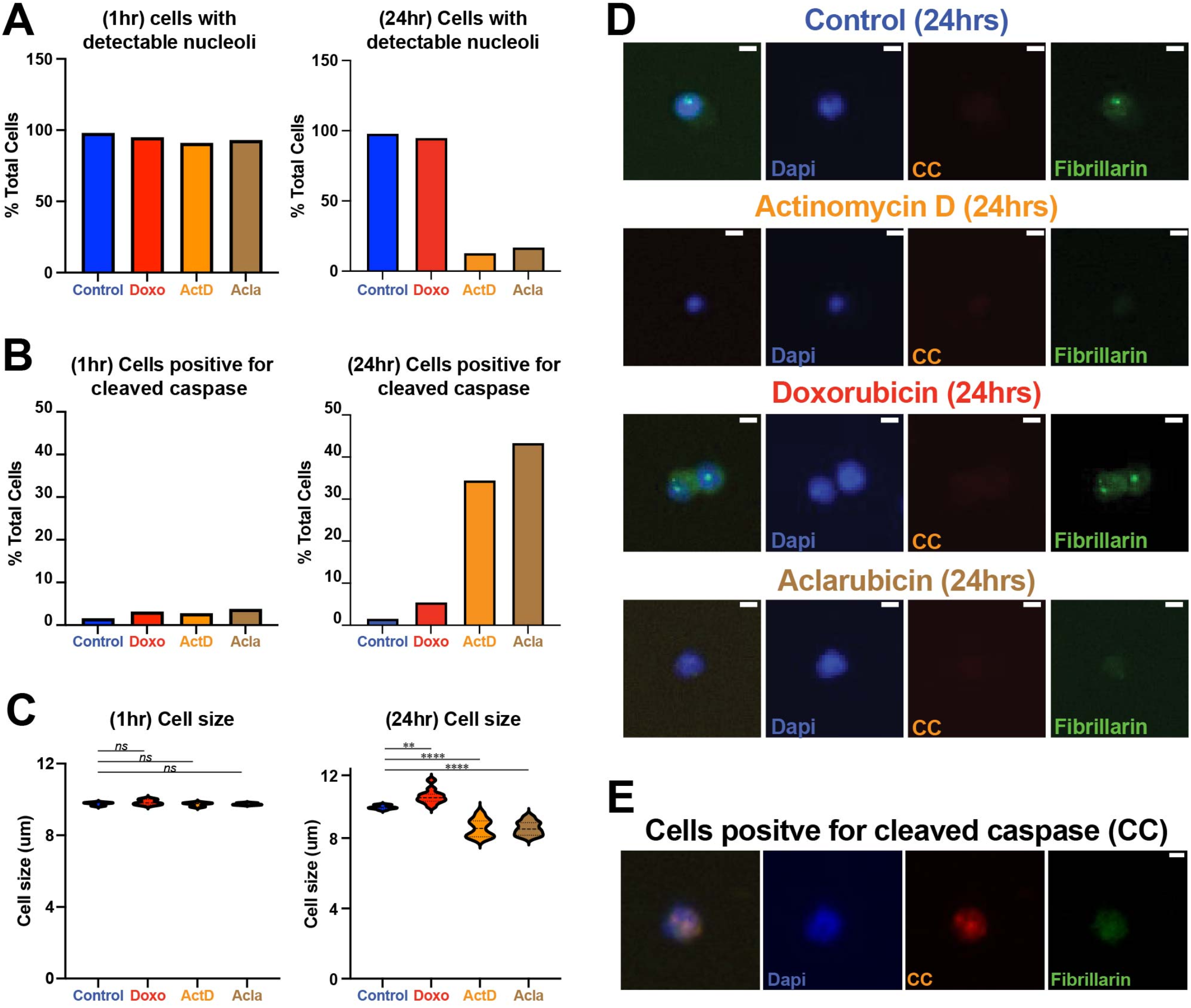
Impacts of drug treatment after 1 hour and 24 hours. (**A**) Percent of cells with detectable nucleolus after 1 hour of treatment and 24 hours treatment. (**B**) Percent of cells with detectable cleaved-caspase (cell death) staining after 1 hour of treatment and 24 hours treatment (**C**) Cell size measurements after 1 hr of treatment and 24hrs treatment. (**D**) Immunofluorescent images of DAPI, Cleaved-caspase (CC) and Fibrillarin (nucleolar marker) after 24 hours treatment. (**E**) Example of cleaved caspase-positive cell. Scale bar = 5μm(A) Immunofluorescent images of DAPI, Cleaved-caspase (CC) and Fibrillarin (nucleolar marker) for cells at 24 hr drug treatment. Scale bar = 5μm. *ns* = not significant, **P<0.01; **** P<0.0001. Ordinary one-way ANOVA with multiple comparisons.

**Fig. S2:**
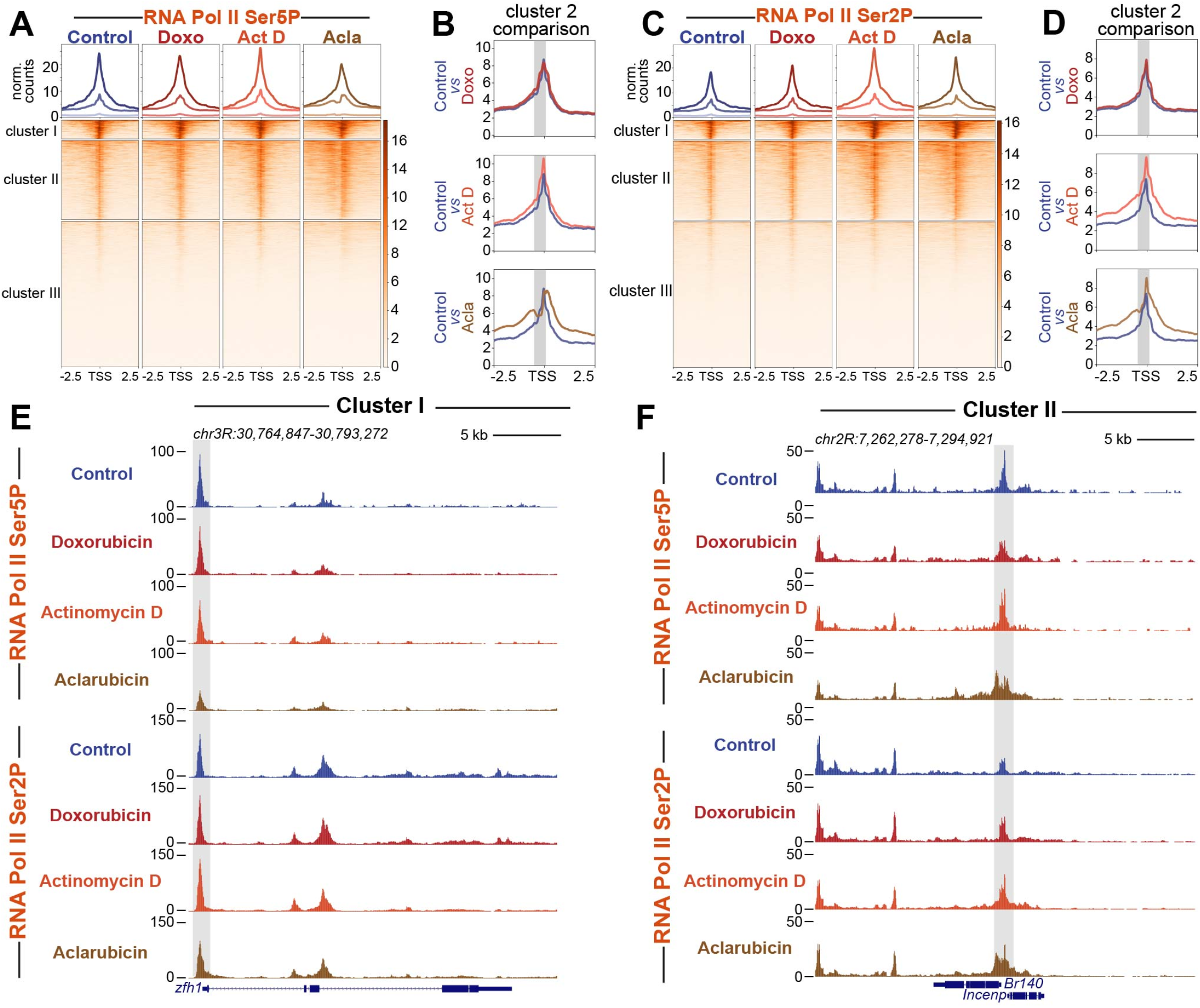
Drug treatment impacts the relative distribution of RNA Pol II Ser5P and Ser2P. (**A**) Heatmap aligned to the transcriptional start site (TSS) of all promoters showing normalized counts of RNA Pol II ser5P CUT&Tag signal clustered via k-means clustering derived from CUTAC datasets (k=3). (**B**) Enlarged comparison of accessibility differences between different drug groups and controls. Gray box marks upstream promoter region. (**C**) Heatmap aligned to TSS of all promoters showing normalized counts of RNA Pol II Ser2P CUT&Tag signal clustered via k-means clustering (k=3) derived from CUTAC datasets. (**D**) Enlarged comparison of G-quadruplex differences between different drug groups and controls. Gray box marks upstream promoter region. (**E**) Representative UCSC browser track snapshot of RNA Pol II ser5P and RNA Pol II Ser2P distribution at Cluster I gene. (**F**) Representative UCSC browser track snapshot of RNA Pol II ser5P and RNA Pol II Ser2P distribution at Cluster II gene.

**Fig. S3:**
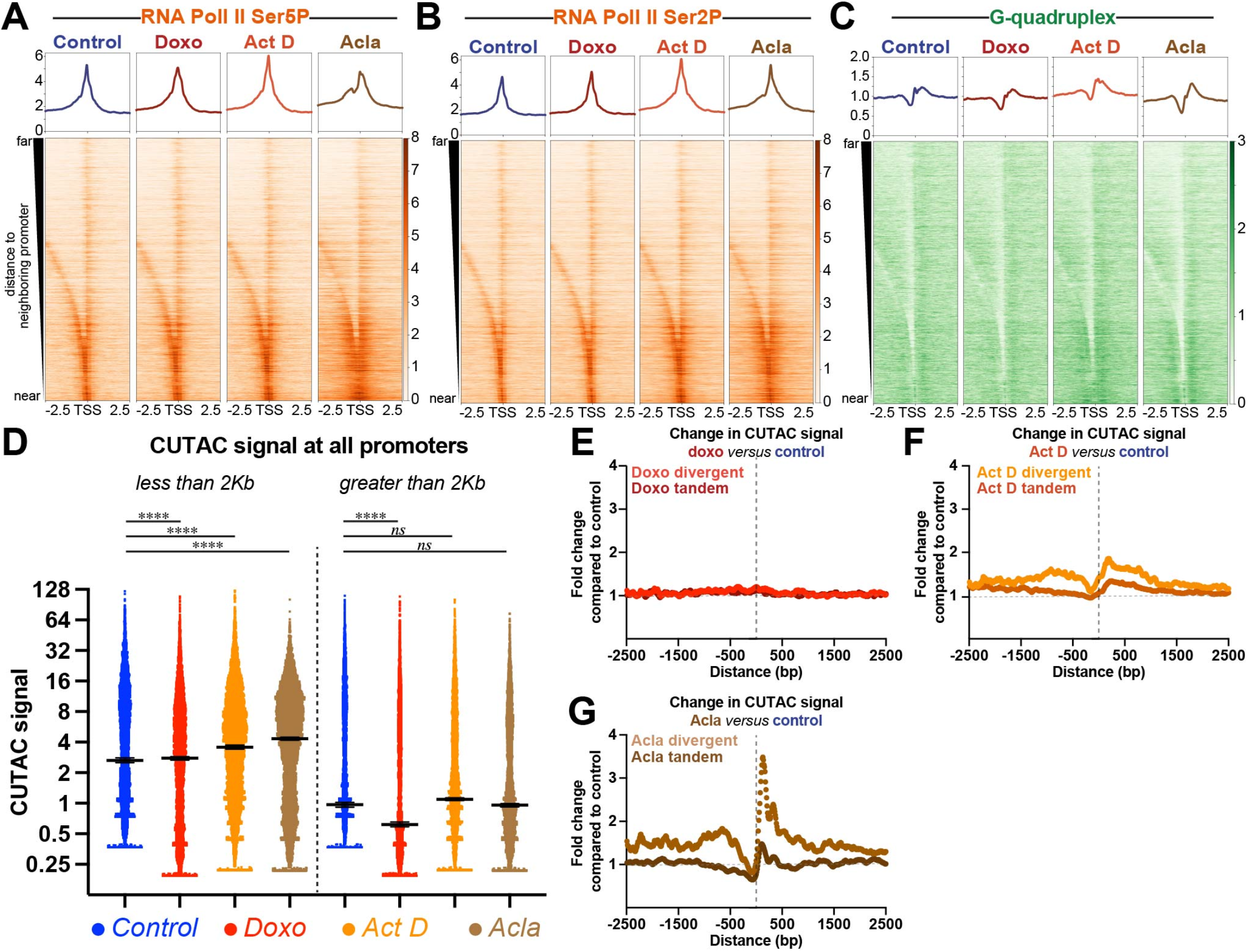
Promoter distance and orientation effects RNA Pol II, G-quadruplexes, accessibility, and torsion. Heatmaps of CUT&Tag data targeting RNA Pol II ser5P (**A**) RNA Pol II ser2P (**B**) and G-quadruplexes (**C**) aligned to the transcriptional start site (TSS) of all promoters sorted by distance to nearest upstream promoter element and plotted in descending order. (**D**) Plot showing CUTAC chromatin accessibility signal for promoters less than 2kb apart and greater than 2kb apart. Median shown as black bar with 95% confidence interval. (**E**) Plot showing fold change of CUTAC chromatin accessibility centered on the TSS for doxorubicin *vs* control (E), actinomycin D *vs* control (**F**) and aclarubicin *vs* control (**G**) Horizontal dotted line indicates 1-fold. Vertical dotted line indicates TSS. (**H**) Plot showing TMP-seq signal for promoters less than 2kb apart and greater than 2kb apart. **** = p< 0.0001 Kruskal-Wallis test with multiple comparisons for panel D.

**Fig. S4:**
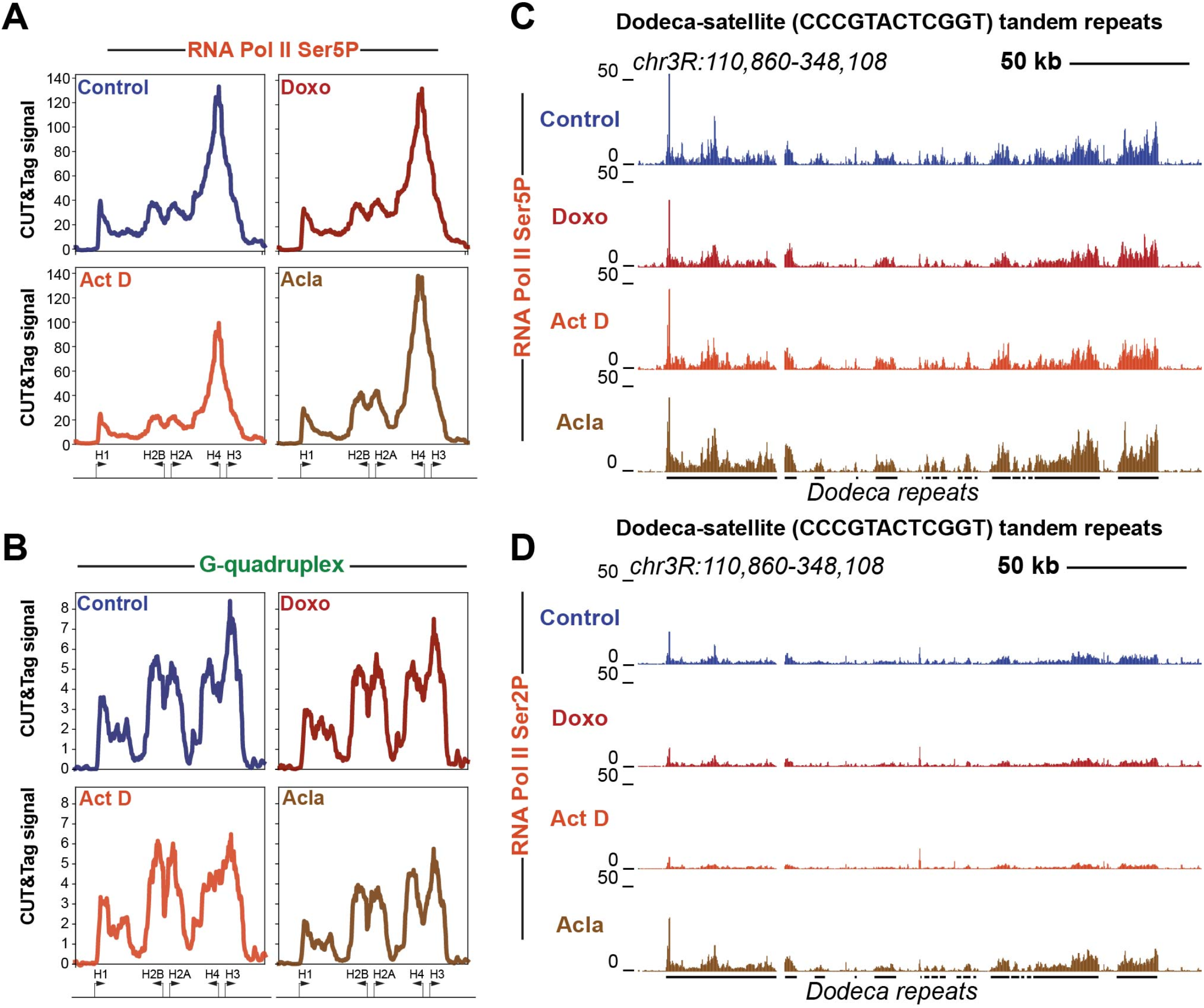
Histone cluster and Dodeca-satellite repeats show distinct responses to drug treatment. (**A**) Average coverage plot of histone clusters showing CUT&Tag data targeting RNA Pol II ser5P. (**B**) Average coverage plot of histone clusters showing CUT&Tag data targeting G-quadruplexes. Arrows at the bottom of average plots indicate approximate positions of histone genes. (**C**) UCSC browser track snapshot of CUT&Tag data targeting RNA Pol II ser5P at Dodeca-satellite repeats. (**D**) UCSC browser track snapshot of CUT&Tag data targeting RNA Pol II ser2P at Dodeca-satellite repeats. Black lines below the browser tracks indicate location of Dodeca-satellite repeats.

## Notes

### Competing Interest Statement

The authors have declared no competing interest.

